# Evaluation of intracochlear pressure and distribution during fluid application in cochlea models and human petrous bone

**DOI:** 10.1101/2024.03.25.586533

**Authors:** R. Kim, C. Riemann, A. Kilgué, S. Schleyer, CJ Pfeiffer, LU. Scholtz, M. Schürmann, I. Todt

## Abstract

**Introduction:** The important factor during the application of substances for an inner ear therapy is the atraumatic execution as well as a homogeneous distribution over the cochlea in a reasonable time frame. Since faster delivery can be obtained with higher pressure but, higher pressure will lead to traumatic execution, there are certain constrains for the delivery process. Because of this, an optimized procedure for intracochlear delivery is needed, which enables reduction of intracochlear pressure during perfusion in reasonable time.

Hence, the aim of this study was to compare different techniques of substance application and their effects on intracochlear pressure in different models.

**Material and Methods:** Intracochlear pressure was measured by fiberoptic pressure sensors in artificial cochlea models and in a human temporal bone. The pressure sensor was introduced into the inner ear models via an additional channel or the lateral arcade of the temporal bone.

In all models the substance was applied by means of an inner ear catheter (MED-EL, Innsbruck, Austria) via the round window with methylene blue with or without second access to the cochlea (helicotrema/oval window).

**Results:** The application of substances showed significant differences in intracochlear pressure and substance distribution at the same velocity between the models with and without second access.

**Conclusion:** Using a second-hole technique leads to a faster homogeneous distribution, as well as a lower intracochlear pressure, which can be assumed to be an essential factor for hearing preservation during substance application.

## 1. Introduction

According to the WHO (World Health Organization), more than 5% of the worlds’ population suffers from hearing loss, hampering participation and hence requiring rehabilitation (1). Systemic therapy or intratympanic local drug delivery, which is clinically used for sudden sensorineural hearing loss, have pharmacokinetic limitations since blood-labyrinth barrier blocks and limits diffusion through the RWM (round window membrane) (2). Therefore, a more directed technique to deliver therapeutics effectively to the inner ear is needed. Considering the advances in the comprehension of the various inner ear pathologies, therapeutic approaches with diverse medications will arise in the near future. Besides the traditional pharmaceuticals based on small molecules with poor but acceptable diffusion potential, newly arisen therapies based on, e.g. gene or cell therapy(3-5), require delivery directly to the inner ear to unlock their potential (6, 7).

This direct administration of a substance was investigated by several research groups, introducing liquid formulations directly into the cochlea. The design of those experiments differs in terms of approach, material or sampling method. In this sense, injections into the perilymph have been performed via needles (8, 9), glass pipettes (10), cochlear catheters (CC) (11) and even during the cochlear implantation (12, 13). With intracochlear injection, the distribution of substance along the scala tympani was effective. However, fluid leakage at the injection site and efflux of fluid at the perforation site were shown to be challenging (14). To prevent leakage, Plontke et al. have used 1% sodium hyaluronate or 17% poloxamer 407 as internal sealing and medical grade biologically-compatible cyanoacrylate adhesive as external sealing at the injection side, which ensured a higher concentration of substance in the perilymph (15). But the problem of a basal-apical concentration gradient or homogeneous distribution has not yet been explained.

During an intracochlear intervention, the intracochlear pressure (ICP) is assumed to be a central factor for the preservation of residual hearing (16, 17). Despite these points, the intracochlear pressure should without any doubt be rising in the closed setting described above as fluids during a cochlear intervention. Furthermore, intracochlear pressure changes during the procedure should be minimized for the atraumatic intervention (18).

To verify substance distribution in the cochlea, sampling is performed at a different location in the cochlea than the application site. It seems obvious that this second perforation for sampling will lead to an improvement of substance distribution. Different groups used the RW (round window) or the lateral semicircular canal, for the application. Sampling was performed in the apex, or oval window (19-21, 11). Groups have observed a basal-apical concentration gradient along the scala tympany, but according to Yildiz et al., a homogenous intracochlear distribution of substance was achieved by using CC (11) at the round window and taking the sample at the oval window.

Despite improved knowledge regarding intracochlear substance distribution upon application, there continues to be an absence of data concerning intracochlear pressure potentially injuring residual hearing. The aim of this study was to compare different techniques of substance application and their effects on intracochlear pressure and substance distribution in different models. The applied methods were chosen to be transferable into clinical practice (timeframe, surgical approach, etc.).

## 2. Material and Methods

### 2.1. General experimental procedure

These two measurements during substance delivery experiments in this study were visual measurements of substance distribution and pressure analysis. For the intracochlear injection, an infusion syringe pump (Fresenius, Bad Homburg vor der Höhe, Germany) ensuring continuous substance application was utilized, this prevented any sound pressure a peristaltic pump would introduce. The experiments were performed in two different types of artificial cochlea models (an unfolded cochlea model and a cochlea model) and in a human petrous bone. Substance distribution was evaluated via photographic images, and intracochlear pressure was measured using fiberoptic pressure sensors (FOP M200, FISO, Quebec, Canada). In all models, substance application was performed using an Inner Ear Catheter (MED-EL, Innsbruck, Austria) via the round window the applied liquid was methylene blue (Merck KGaA, Darmstadt, Germany)-stained saline (0.9 %). The configurations of the model were with and without second access to the cochlea (helicotrema / oval window). Pressure measurements were performed via an additional burr channel of the lateral semicircular canal. Perforation for the connection of a catheter and pressure sensor as well as for the creation of second access was performed by a 0.4 mm perforator (Karl Storz, Tuttlingen, Germany). Measurements were performed at the time of 0, 210, 270, 390, 450, 510, 630, 750 and 900 seconds after the onset of substance perfusion.

### 2.2. Models

Two 3D-printed artificial models were utilized during experiments. The first model was a linear model, representing an unfolded cochlea with a length of 3 cm and a volume of 100 μm^3^, which is anatomically in the average range of a cochlea (22). It consists of a cone like hollow space and a diameter of without any anatomical structure such as the scala tympani, scala media or scala vestibuli. It has an additional drilled hole on the apex (Figure. 1A).

**Fig 1.**
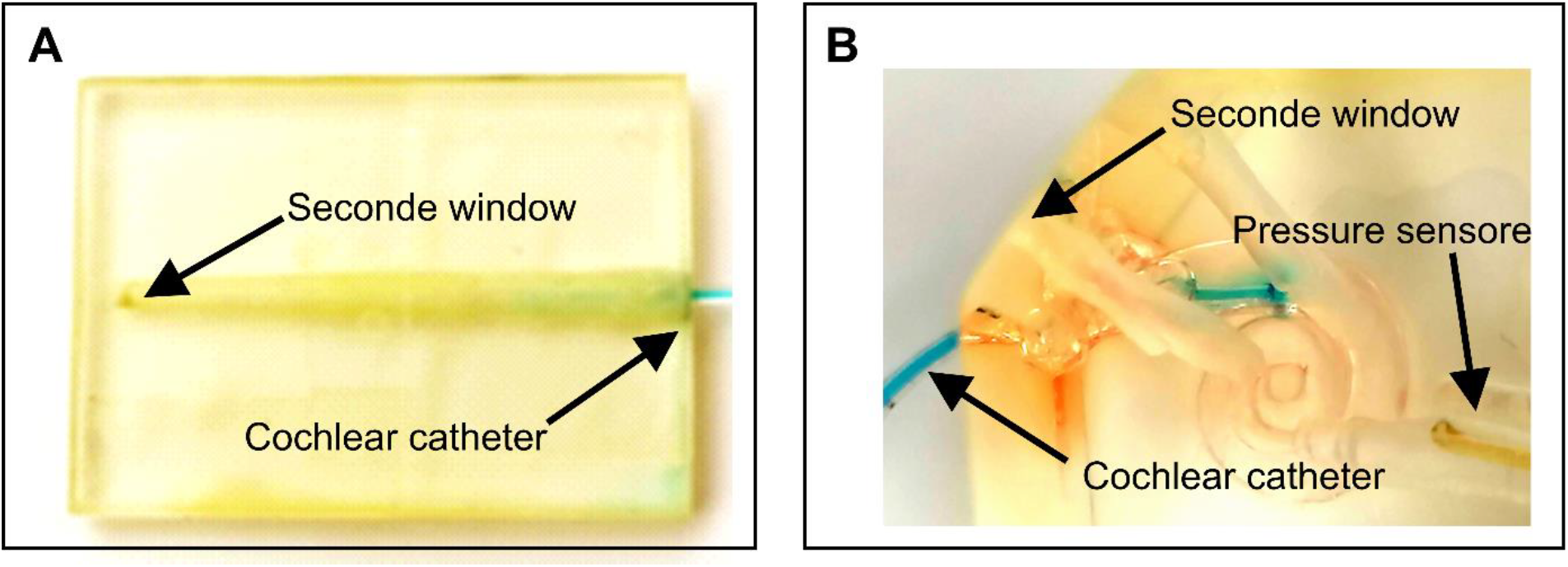
(A) The linear model (B) The cochlea model, Cochlear catheter inserted.

The second model is a full-scale model of the cochlea with a volume of 87 mm^3^. The artificial cochlea model has the same shape and size as a human cochlea but without any separation of anatomical structure, such as the scala tympani, scala media or scala vestibuli. On the apex and also on the middle of the cochlea, there are several additional burr channels (Figure. 1B). In contrast to linear model, the cochlea model has two more drilled holes, one for the pressure sensor and the other serving as second window (Figure 1B).

### 2.3. Cochlear Catheter

The cochlear catheter used in our study is silicone-based Inner Ear Catheter for the intracochlear administration of substances (MED-EL, Innsbruck, Austria) (23). Black dots on the top of the catheter down mark the 5 mm, 10 mm, and 15 mm respectively. The reservoir at the posterior end is punctured with a needle to enable loading with the test substance.

In our experiments, we have inserted the substance-filled catheter up to 5 mm in to RWM.

### 2.4. Pressure Sensor

The ICP was measured using a microoptical pressure sensor FOP M200 (FISO, Quebec, Canada) with a tip size of 0.2 mm. The tip of the pressure sensor is a hollow glass tube sealed on one end by a thin plastic film diaphragm coated with a reflective surface of evaporated gold. A optical fiber is located within the glass tube at a small distance (50–100 μm) from the diaphragm tip. The optical fiber is attached to a LED light source and a photodiode sensor. Light from the LED source reaches the sensor tip of the optical fiber, the beam widens as it exits the fiber and is reflected by the gold-covered flexible diaphragm. The reflected light is sensed by the photodiode. Small distance displacements of the diaphragm modulate the intensity of reflected light. The sensor is connected to a module that is linked to a computer. Evolution software (FISO, Quebec, Canada) was used to record the ICP. The time resolution of the sensor was set to 300 Hz, leading to an accuracy of 3 %.

### 2.5. Fibrin glue

To avoid the leakage at the punctured site, we used fibrin glue, Tisseel (Baxter international, Deerfield, USA), after the insertion of the cochlear catheter and pressure sensor. At each application, approximately 1 ml of fibrin glue was used. To ensure complete curing, we waited at least 2 minutes after sealing before the measurements were started.

### 2.6. Experimental setup

#### 2.6.1. Substance injection in the linear Model

Prior to the substance application, the empty space in the cochlea model, including the drilled hole, was completely filled with saline (0.9 %). The drilled hole and the opening on the base were covered with tape. Afterwards, the aperture on the base was punctured with a 0.4 mm perforator, and the cochlear catheter was inserted up to the first black marking at 5mm. In one setting, ca. 1 ml of fibrin glue was applied around the opening after Insertion of CC to avoid leakage. The experiment was performed with and without a second hole. The second hole was generated at the apex. After 2 minutes, the substance could be injected into the cochlea continuously per syringe pump under 2 ml/h and 4 ml/h. The distribution was captured photographically at 0, 210, 270, 390, 450, 510, 630, 750 and 900 seconds after the beginning of substance application. The experiments were performed in series with a CC in an unchanged position to exclude CC position-related bias and to allow interexperimental comparability.

#### 2.6.2. Substance injection in the cochlea model

The puncturing and insertion of CC is performed identically as in the linear model (see 2.6.1.). A pressure sensor is also inserted in the burr channel of the cochlea. To avoid leakage, ca. 1 ml fibrin glue was applied around the opening. The experiment was performed with and without a second hole. The second hole was generated at the apex. After fibrin glue application and an additional 2 minutes pause, the substance was continuously injected in the cochlea by a syringe pump at a rate of 2 ml/h and 4 ml/h. The distribution was captured photographically, and the pressure measurement could be checked at 0, 210, 270, 390, 450, 510, 630, 750 and 900 seconds after the beginning of substance application.

#### 2.6.3. Substance injection in the human petrous bone

In the human petrous bone, we measured intracochlear pressure during substance application. First, we opened the round window, and punctured the lateral semicircular canal (LSC) with a 0.4 mm perforator, and filled it with fluid. In the RW a substance-filled cochlear catheter was introduced and a pressure sensor was inserted into the window of the LSC. Afterwards, we covered both sites with connective tissue and fibrin glue. The second hole was created with a 4 mm perforator in the stapes footplate. The pressure measurement with substance administration was performed at an application rate of 0.4 ml/h for 15 minutes. The evaluation of intracochlear pressure values was carried out at the same time periods as the previous experiments.

### 2.7. Analysis

For the measurement of substance distribution in our linear cochlea model, the ratio is calculated as shown at figure 3A. For the cochlea model, the distribution was determined by measuring the distribution angle reached, as shown in the figure 3B. The intracochlear pressure was measured continuously and the value was measured with the Evolution software (see Material and Methods).

**Fig 2.**
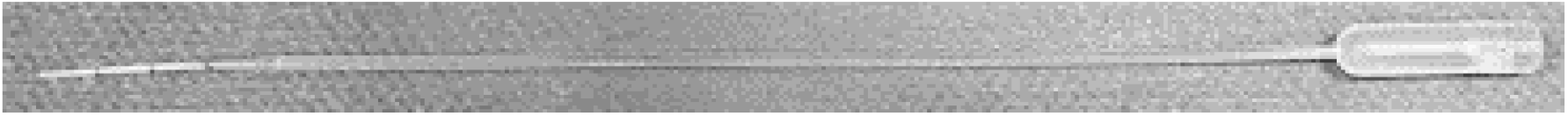
cochlear catheter.

**Fig 3.**
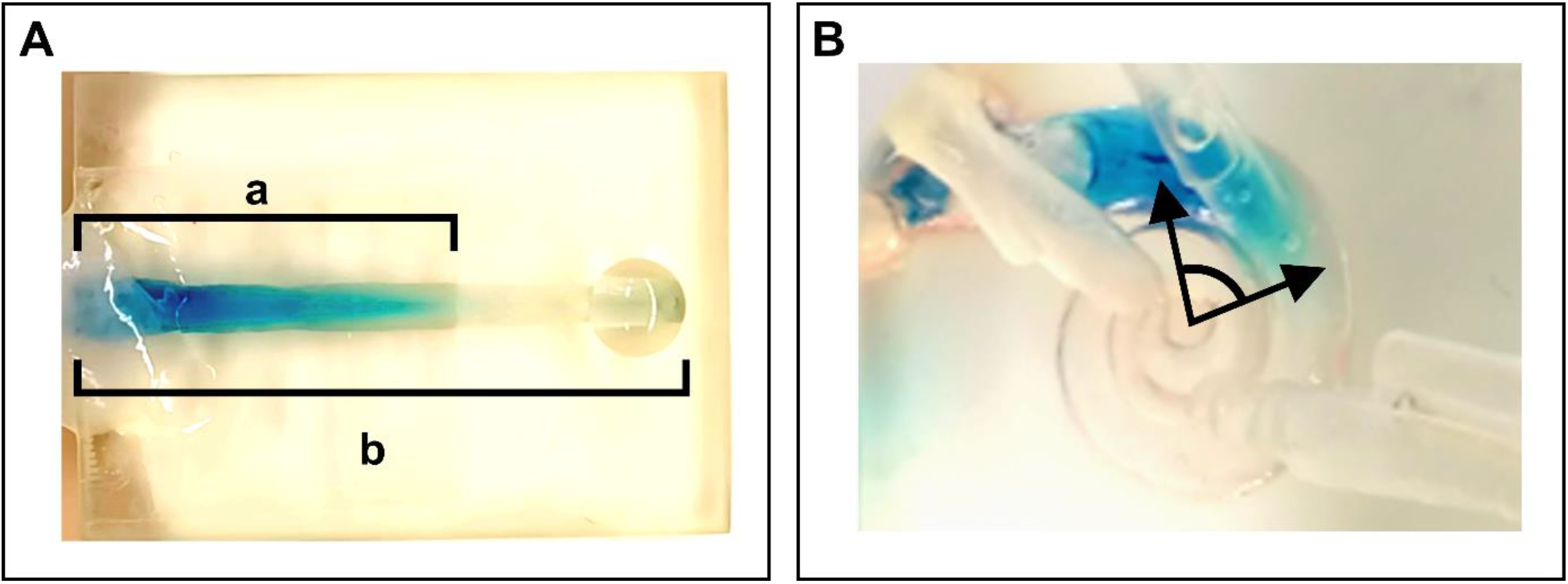
Measurement approaches in the two models. **(A)** measurement of substance distribution in the linear model as a ratio a/b. **(B)** measurement of substance distribution in the cochlea model as an angle

## 3. Results

### 3.1. Substance distribution with or without second hole and without sealing

In order to determine whether the second-hole technique allows a homogeneous fluid distribution during intracochlear injection and also to confirm if sealing after puncturing affects the fluid distribution, we first performed the experiment in two different models without sealing.

In the linear model, a maximal distribution ratio of around 0.4 could be reached after 15 minutes for all conditions. While the puncturing seems to have increased this final ratio at 0.2 ml/h up to 0.6, this difference was not significant (Figure 4A).

**Fig 4.**
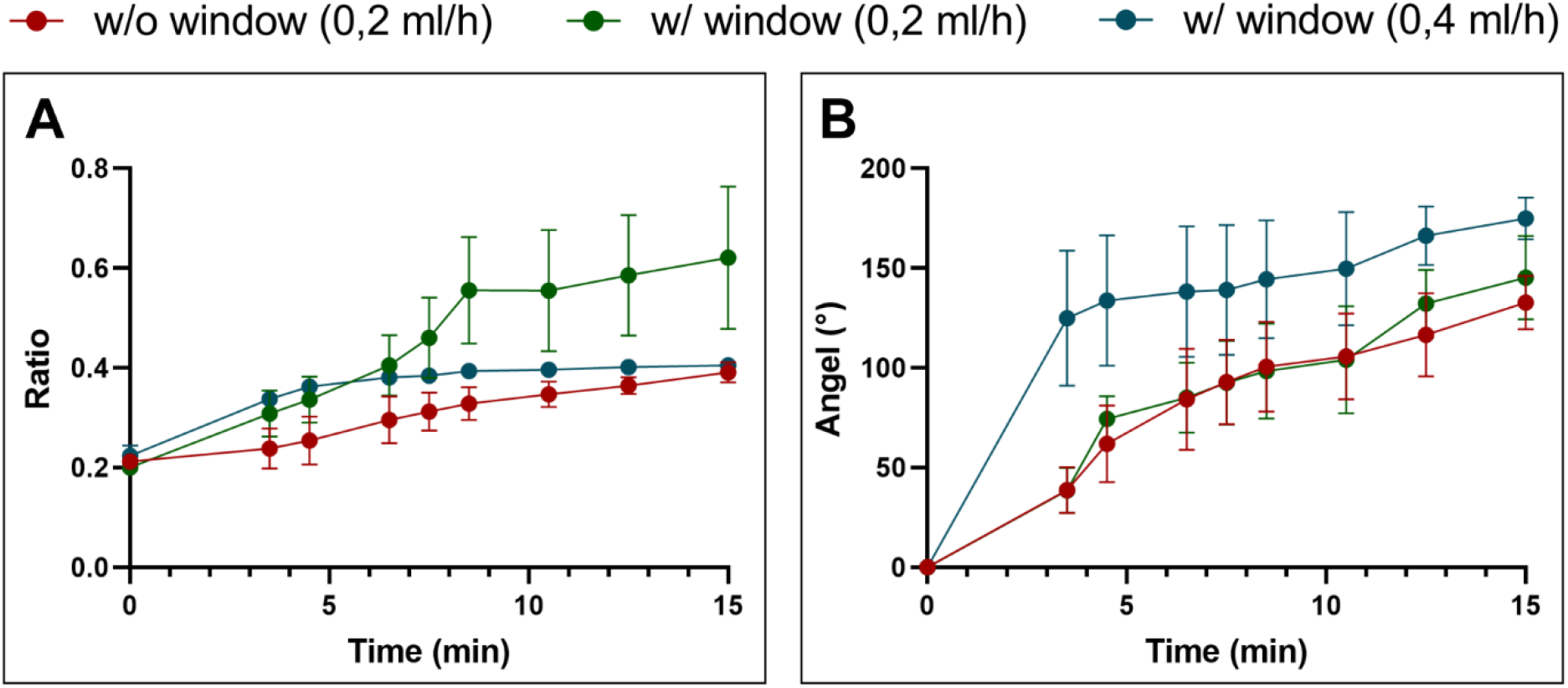
Measurements of the distribution of fluid in the linear and cochlea models without fibrin sealant and with and without a second window. **(A)** In the linear model, a maximal distribution ratio of around 0.4 can be reached after 15 minutes. The puncturing seems to increase this final ratio to 0.2 ml/h, but this difference is not significant. **(B)** In the cochlea model, a poor distribution of the test fluid upon omission of fibrin sealant is observed. The angle of distribution only reaches about 140° after 15 minutes of perfusion. Likewise to the linear model, the puncturing seems to improve the perfusion, but without the difference reaching a level of significance. The data are presented as a mean ratio ± standard deviation.

In the cochlea model, a poor distribution of the test fluid without fibrin sealant was also observed. The angle of distribution merely reached about 133° without second hole and 145° with second hole at 0.2 ml/h after 15 minutes of perfusion. Under higher perfusion rate at 0.4 ml/h with second hole, the distribution was reached about 175°. Similar to the linear model, puncturing seems to improve the perfusion, but without the difference reaching a level of significance (Figure 4B).

Figure 5 shows the leakage as the experiment was performed without sealing. A leakage on the insertion site affects the perfusion and occurs with a non-homogenous distribution.

**Fig 5.**
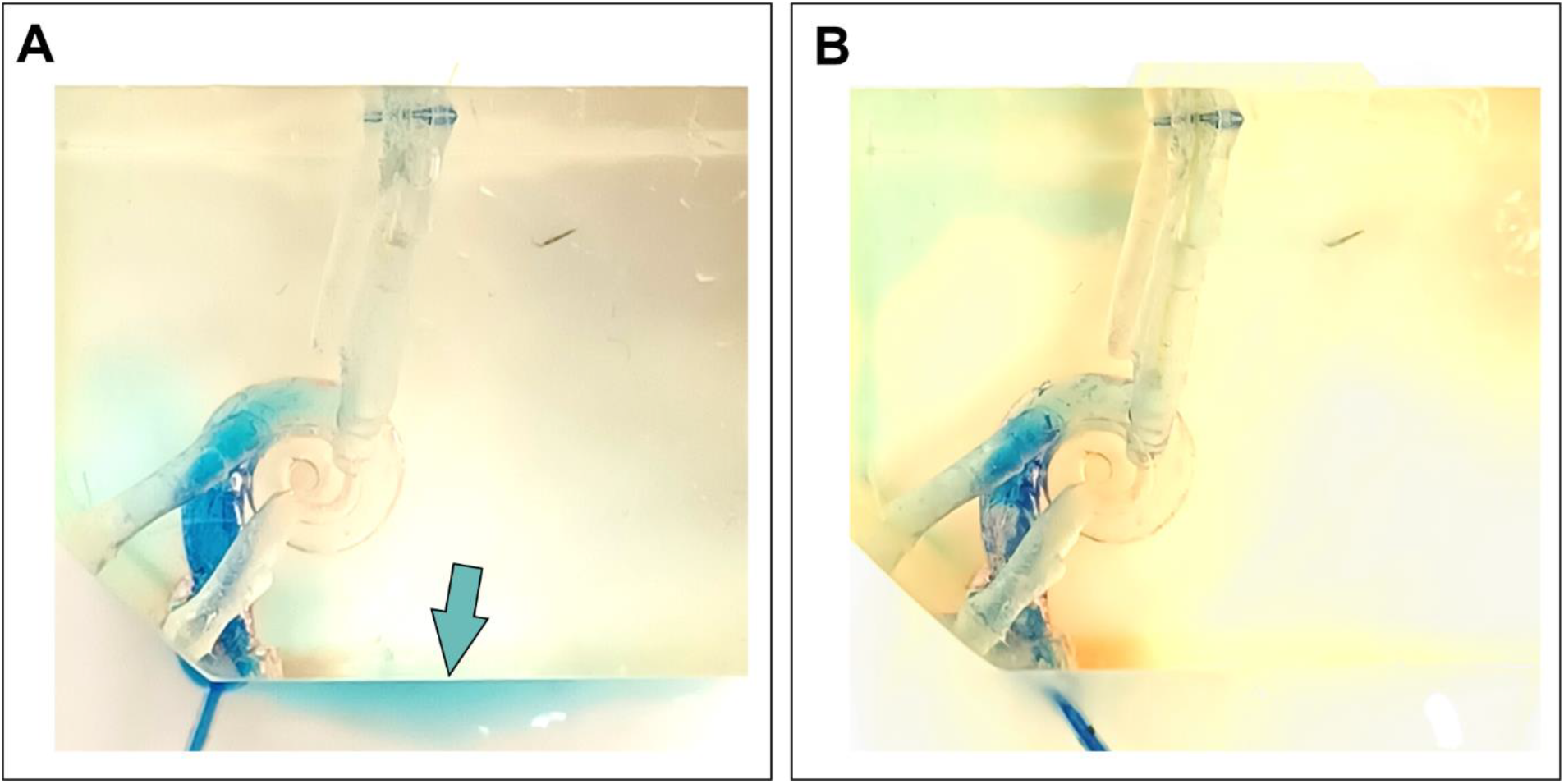
Lag of the fibrin sealant results in leakage from the site of insertion in the cochlea model. **(A)** The perfusion of the cochlea model without fibrin sealant results in a distinct spillage of the test fluid after the onset of perfusion (blue arrow). **(B)** The sealing of the perfusion site prevents any spillage of the test fluid and leads to a clean perfusion of the cochlea model.

### 3.2. Substance distribution with or without a second hole and with sealing

The fluid distribution after sealing the intracochlear access of the CC via RWM was more homogenous and faster than without a sealing procedure. In the linear model, the sealing of the site of perfusion in combination with the higher perfusion rate of 0.4 ml/h leads to a complete distribution of the test liquid along the whole cochlea model after about 7.5 minutes. The substance application with an additional second window at a rate of 0.2 ml/h could lead to better perfusion than without a second window, but not to a complete distribution in the linear model (Ratio 0.31 vs. 0.54). At the end of experiment we observed the significant difference of perfusion between with or without second hole at a rate of 0.2 ml/h, but also with different perfusion rate between 0.2 ml/h and 0.4 ml/h as the second hole perforated. In addition, the sealing leads to a higher reproducibility of the measurements, which results in a significant discrimination between the three experimental conditions (Figure 6A).

**Fig 6.**
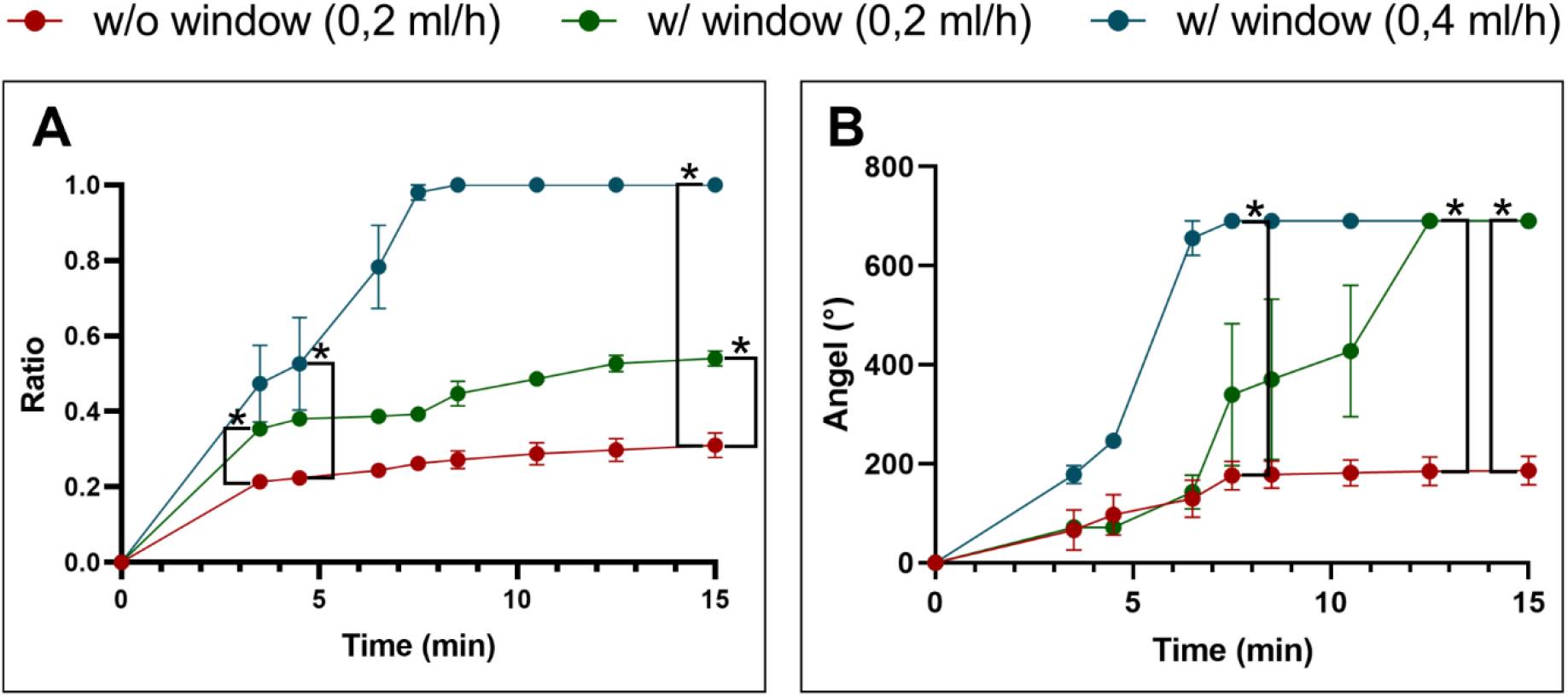
Measurements of the distribution of test liquid in artificial models with fibrin sealant and with and without puncturing. **(A)** In the linear model, the sealing of the site of perfusion in combination with the relative high perfusion velocity of 0.4 ml/h leads to a complete distribution of the test liquid along the whole cochlea model after about 8.5 minutes. In addition, the sealing leads to a higher reproducibility of the measurements, which results in a significant discrimination between the three experimental conditions. **(B)** In the cochlea model, the fibrin sealing results in a complete perfusion of the cochlea model with the test fluid at all perfusion velocities, but only in the models with a second window. The distribution of the test liquid can be distinguished significantly between these two models after the establishment of a complete perfusion. This occurred after 7.5 minutes. at 0.4 ml/h and after 12.5 minutes at 0.2 ml/h (Mann-Whitney-U test, two-tailed, confidence interval 95%, *=p<0.05).

In the cochlea model, the fibrin sealing results in a complete distribution of the test fluid at all perfusion velocities in the cochlea model with a second window. The distribution of the test liquid differs significantly between the investigated perfusion velocities after the establishment of a complete perfusion. This occurred after 8.5 minutes at 0.4 ml/h and after 12.5 minutes at 0.2 ml/h (Figure 6B).

### 3.3. Intracochlear pressure in the cochlea model

The pressure changes in the cochlea models were measured at certain time points (Figure 7) continuously. Each experimental condition was measured at least three times. The pressure inside the model with a second window were lower and more constant than the intracochlear pressure in models without a second window. With a second window, the intracochlear pressure is almost constant around 0 mmHg. The intracochlear pressure of the models without a second window increased continuously. This leads to significant differences between the different perfusion velocities after 7.5 minutes.

**Fig 7.**
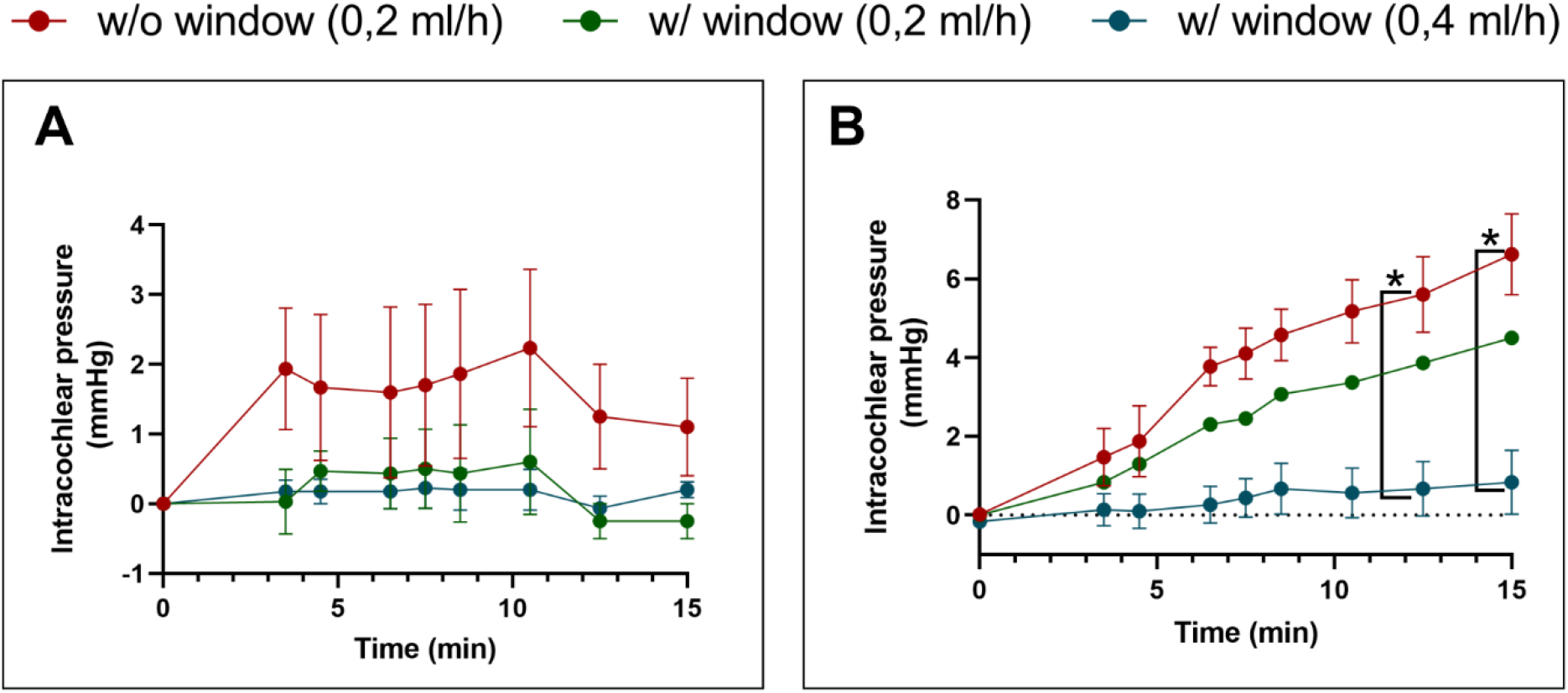
The pressure changes in the cochlea model during perfusion with and without puncturing or fibrin sealant. **(A)** The pressure changes in the cochlea model without fibrin sealing. The pressure inside the models with puncturing are distributed around 0 mmHg. The average value for the model without puncturing is between 1 and 2 mmHg. At the end, the difference between these two models does not reach the level of statistical significance. **(B)** The intracochlear pressure of the models sealed with fibrin. Likewise, with the models without fibrin sealant, the pressure stays close to 0 mmHg with perfusion at a rate of 0.4 ml/h. A much steeper increase can be observed at 0.2 ml/h with or without puncturing. This leads to significant differences between the different perfusion velocities after 7.5 minutes. (Mann-Whitney-U test, two-tailed, confidence interval 95%, *=p<0.05)

**Fig 8.**
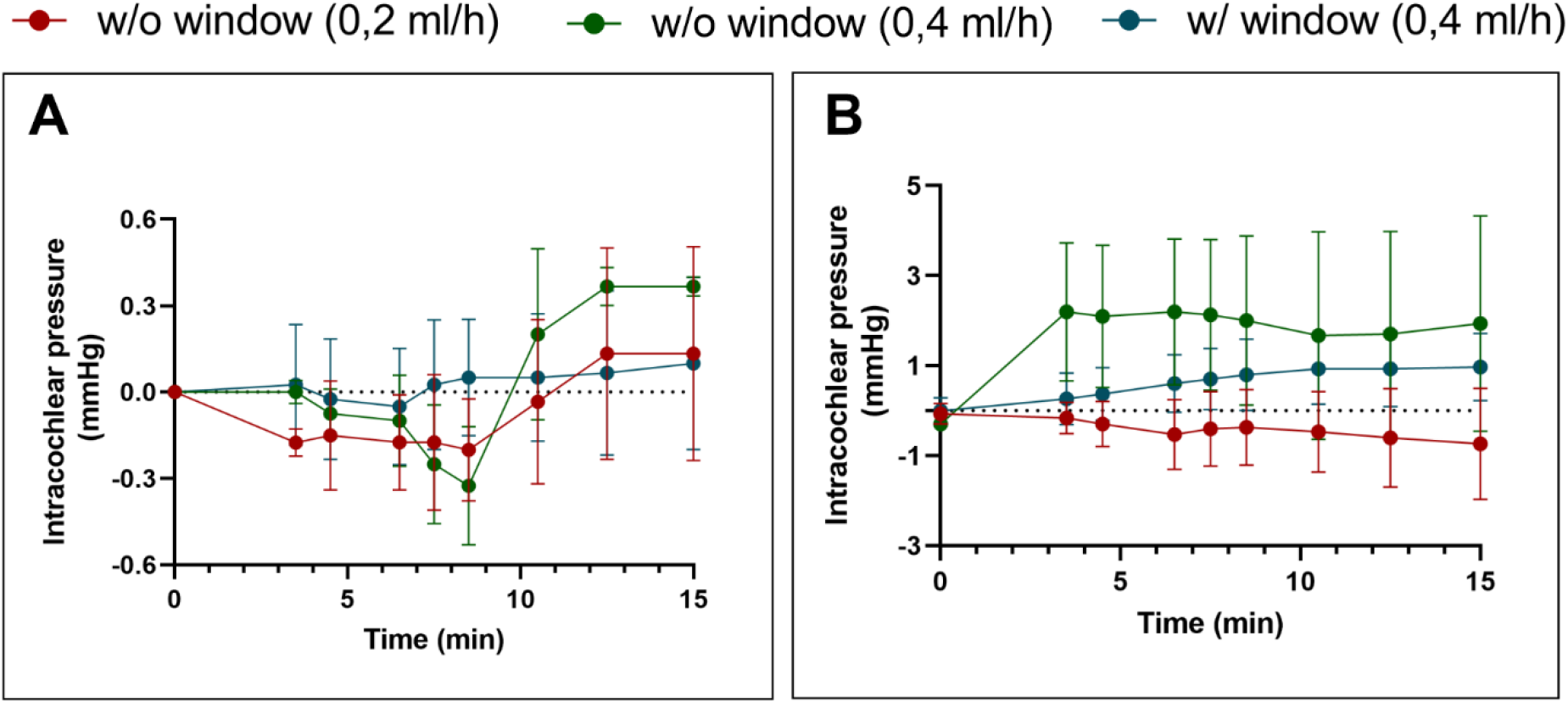
Measurements of the intracochlear pressure in the mastoid bone model, with and without puncturing or fibrin. **(A)** Without sealing the site of perfusion, the pressure fluctuates near 0 mmHg. Due to a relatively large deviation, no difference between the conditions can be assessed. **(B)** Displays the same model but with a sealed perfusion site. It can be assumed that the sealing leads to a higher pressure than 0 mmHg for the model with puncturing and a perfusion rate of 0.2 ml/h. But due to the large deviations between the technical replicates, no assumption can be made whether there is a real difference between these single conditions.

### 3.4. Intracochlear pressure in the human petrous bone

Performing the application of the substance with a second window, the observed pressure fluctuates near 0 mmHg with a rate at 0.4 ml/h without sealing the site of perfusion in compare to the experiments without a second window and showed the distinct fluctuation range of intracochlear pressure without significance (0.18 vs. 0,73 mmHg). The intracochlear pressures were not stable as well as with a fluid application rate at 0.2 ml/h without second hole. But the fluctuation range of intracochlear pressure was larger with a rate of 0.4 ml/h than with 0.2 ml/h (0.48 vs. 0.73 mmHg).

As the pressure measurement performed with sealing on the site of perfusion as well as on the insertion site of pressure sensor, the intracochlear pressure were increased continuously with a rate at 0.4 ml/h without and with a second window. But the fluctuation range of intracochlear pressure was larger without second window then with second window (1.63 mmHg vs. 1 mmHg). Without second window the pressure were decreased continuously under 0 mmHg with perfusion at a rate of 0.2 ml/h. During this experiment none of these models reached the level of statistical significance.

The application of substances showed a significant difference in the distribution rate under the same application velocity between the second-hole and single-hole techniques. Furthermore, there was a significant difference in intracochlear pressure between the single-hole technique and the second-hole technique. Additional optimization by application sealant led to a homogenization of the results.

## 4. Discussion

The application of substances (vectors, smart molecules, medication) for inner ear therapy directly via intracochlear is nowadays highly discussed. Because of biological hurdles like the blood labyrinth barrier (BLB), a systemic therapy is hindered, and the poor diffusion prevents efficient intratympanic delivery. Hence, a new delivery approach for inner ear therapy is desperately needed (24).

Important for all substance applications to the inner ear is the need for an atraumatic application. To the best of our knowledge, this is the first study that demonstrates the effect on substance distribution and intracochlear pressure during a second-hole technique for intracochlear substance application. In the cochlea model, we could show that the homogenous substance distribution was possible within 7 minutes, a time frame transferable into a clinical setting.

In the linear model, we observed that the homogenous distribution depends on the application of fibrin glue as well as the rate of injection. In all the experiments performed without using fibrin glue, the substance delivery was incomplete. We think this depends on perforation size or how tightly the CC fitted into the perforation, respectively. Using a fibrin glue increased the reproducibility of the measurements by a remarkable amount. The necessity of sealing after perforation has already been suggested in several studies (15, 14). Plontke et al. described that the utilization of gel or adhesive prevents fluid leakage, and the amount of applied drug or marker in the perilymph was higher compared to that delivered without sealing. Despite the sealing with fibrin glue in our study, we rarely observed the leakage, which could be prevented through second window or by an increased amount of fibrin glue and a sufficient time period for the solidification of fibrin glue. Future studies also need to demonstrate the sealing of the second hole after substance application to prevent perilymph leakage.

The handling of intracochlear pressure is assumed to be an essential factor for hearing preservation during intracochlear procedures (25). In our study, we measured the pressure during substance application in a cochlea model as well as in the human petrous bone. In the cochlea model, the second-hole technique appeared to ensure the stabilization of the intracochlear pressure at different application velocities. Within the human petrous bone model, the intracochlear pressure appeared to stay lower with a second-hole technique at a perfusion rate of 0.4 ml/h. An unexpected result was that under the experiment in a closed setting without a second hole at a perfusion velocity of 2 ml/h, the intracochlear pressure was much lower in the experiment with a second hole at a perfusion velocity of 4 ml/h. We assume that the fluid resorption of dried petrous bone was responsible for this phenomenon. The applied fluid was resorbed in dried bone, and it could exert a negative pressure in the closed setting of the cochlea. It remains to be demonstrated whether the experiment in a humid human petrous bone, which provides the most similar microstructure of the human in vivo cochlea, would show the similar results.

In our study, the measurement of substance distribution was performed through visual evaluation. Previously, study groups, that evaluated the intracochlear substance injection and distribution evaluated the distribution by the concentration of makers such as polysaccharide fluorescein isothiocyanate dextran (FITC-d) or trimethylphenylammonium (TMPA) (21, 11). In the Future, the measurement through mass spectrometry or gas or liquid chromatograph could deliver more precise information pharmacokinetically.

Since all models were cochlea models without implemented scalar separations, the respective effects on fluid distribution and pressure could not be derived. Therefore, the investigation of similar perfusion settings demonstrated in this study, in cochlea models with other inner ear structures (e.g., scala media, scala vestibuli, scala tympani) would be of interest. Furthermore, general limitation e.g. the mechanical properties of the utilized artificial models compared to that of inner ear tissue might confine the transferability of the results obtained with our model into the in vivo situation.

The goal of this study was to suggest procedures facilitating drug, smart-molecules, gene or cell-based therapies for the inner ear via intracochlear application. Hearing preservation, homogenous substance distribution, preventing complications like leakage, and the transferability of procedures into a clinical timeframe are essential aspects of such techniques. The second-hole technique could fulfill those criteria.

## 5. Conclusion

Using a second-hole technique leads to a faster homogeneous distribution, as well as a lower intracochlear pressure, which can be assumed to be an essential factor for hearing preservation during substance application.

